# Pose-gait analysis for cetaceans with biologging tags

**DOI:** 10.1101/2021.12.13.472407

**Authors:** Ding Zhang, Kari Goodbar, Nicole West, Veronique Lesage, Susan Parks, Dave Wiley, Kira Barton, K. Alex Shorter

**Affiliations:** Department of Mechanical Engineering, University of Michigan, Ann Arbor, MI, USA; Dolphin Quest Oahu, Honolulu, HI, USA; Fisheries and Oceans Canada, Canada; Department of Biology, Syracuse University, Syracuse, NY, USA; National Oceanic and Atmospheric Agency’s (NOAA) Stellwagen Bank National Marine Sanctuary, USA

## Abstract

Biologging tags are a key enabling tool for investigating cetacean behavior and locomotion in their natural habitat. Identifying and then parameterizing gait from movement sensor data is critical for these investigations. But how best to characterize gait from tag data remains an open question. Further, the location and orientation of the tag on an animal in the field are variable and can change multiple times during deployment. As a result, the relative orientation of the tag with respect to (wrt) the animal must be determined before a wide variety of further analyses. Currently, custom scripts that involve specific manual heuristics methods tend to be used in the literature. These methods require a level of knowledge and experience that can affect the reliability and repeatability of the analysis. The authors of this work argue that an animal’s gait is composed of a sequence of body poses observed by the tag, demonstrating a specific spatial pattern in the data that can be utilized for different purposes. This work presents an automated data processing pipeline (and software) that takes advantage of the common characteristics of pose and gait of the animal to 1) Identify time instances associated with occurrences of relative motion between the tag and animal; 2) Identify the relative orientation of tag wrt the animal’s body for a given data segment; and 3) Extract gait parameters that are invariant to pose and tag orientation. The authors included biologging tag data from bottlenose dolphins, humpback whales, and beluga whales in this work to validate and demonstrate the approach. Results show that the average relative orientation error of the tag wrt the dolphin’s body after processing was within 11 degrees in roll, pitch, and yaw directions. The average precision and recall for identifying relative tag motion were 0.87 and 0.89, respectively. Examples of the resulting pose and gait analysis demonstrate the potential of this approach to enhance studies that use tag data to investigate movement and behavior. MATLAB source code and data presented in the paper were made available to the public (https://github.com/ding-z/cetacean-pose-gait-analysis.git), with suggestions related to tag data processing practices provided in this paper. The proposed analysis approach will facilitate the use of biologging tags to study cetacean locomotion and behavior.

## Introduction

Biologging tags use a combination of sensors (e.g., accelerometer, magnetometer, gyroscope, speed, pressure sensor, camera, micro/hydrophone) to record data about animal movement, behavior, and the environment. For studies of cetaceans, in particular, many research topics utilize data from biologging tags. For example, tags are commonly used to study the bioacoustics of marine mammals [1–3]. In addition, the biomechanics, time budget, and diving behaviors of cetaceans are also often studied using biologging tags [4–10]. The kinematic data from these tag systems are used to estimate the animal’s position in the environment and provide information about animal behavior in the context of habitat usage [11–18]. Machine learning-based approaches have been applied to tag data to infer behavior when direct human observation/interpretation is expensive, time-consuming, or not possible [19–22]. The key to many of these studies is accurately estimating animal pose (roll, pitch, and yaw, Figure 1).

**Fig 1.**
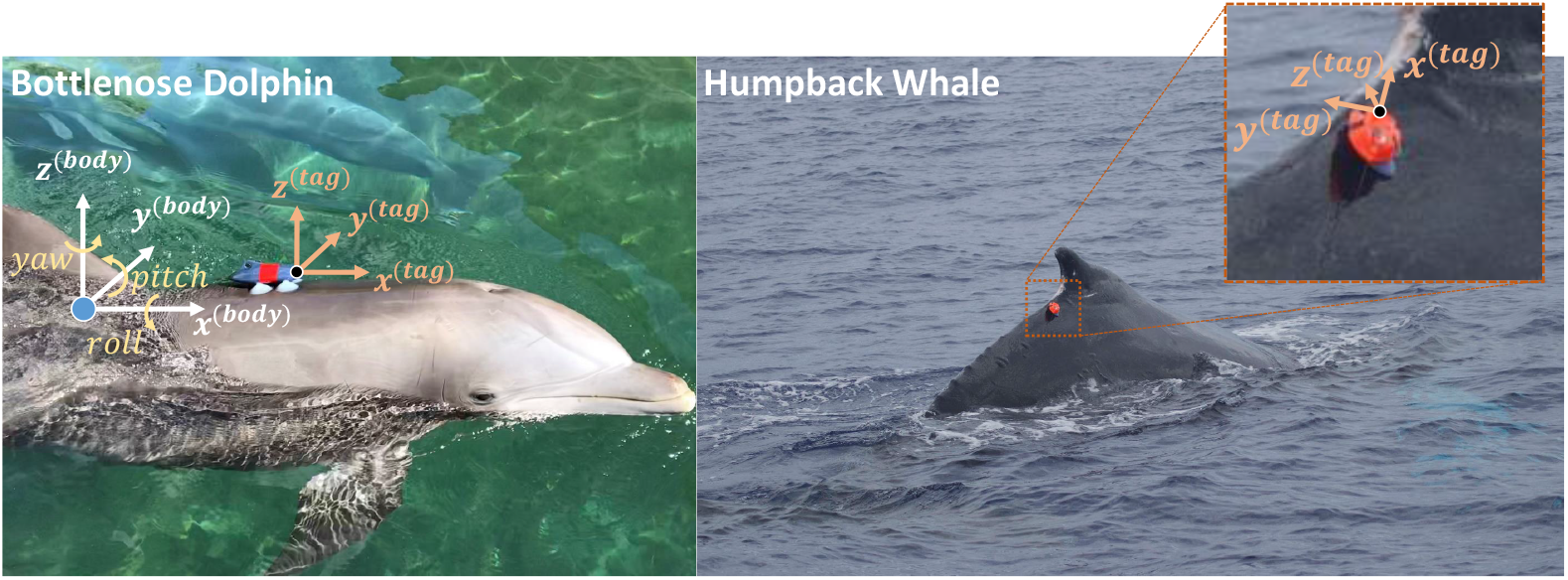
Biologging tag attached to a bottlenose dolphin (MTag, **left**) and a humpback whale (DTag, **right**) with coordinate definitions. Sometimes the tag’s coordinates are aligned with the animal (**left**) but oftentimes they are not (**right**). It is also possible that a tag’s position on the animal changes during deployments. For applications like animal pose estimation, tag data needs to be represented in the animal’s body coordinates during post-processing, which implicitly requires the relative orientation between tag and animal. Note that a positive animal pitch is a negative rotation around the body’s *y*-axis under this coordinate system.

The pose is essential for studies that require an estimated spatial trajectory of the animal or investigate locomotion and gait [4, 5, 8, 11–15, 23]. In the literature, the pose is typically estimated using accelerometer and magnetometer data [1, 8]. When gyroscope data is available, methods like [24] can be used to improve the estimated animal pose. Swimming gait is essentially composed of a repetitive sequence of poses, but how to efficiently parameterize the gait of the animal from a pose sequence remains an open question. Studies in the literature commonly use pitch to identify and quantify gait via parameters like frequency, amplitude, and duty-factor [4, 5, 8]. But these pitch-based gait descriptors are sensitive to the roll angle of the animal. For example, the pitch of an animal derived from accelerometer data captures little to no information about the animal’s gait if the animal is fluking sideways (a 90-degree roll angle). To characterize an animal’s gait, we need a measure wrt the animal’s body rather than the earth’s horizontal plane. To this end, the measured angular rate from a gyroscope can be integrated to estimate the rotation angle wrt the animal itself. However, this estimation is subject to accumulated sensor error, and gyroscopes are unavailable in some tag platforms. As such, it is essential to develop approaches that can address this in accelerometer-based estimates of orientation.

Because many cetacean tags use a suction cup-based attachment method, the location and orientation of the tag wrt the animal can change during field deployments [1, 25]. As a result, the relative orientation of the tag wrt the animal must be determined before animal pose calculation [1, 8, 26]. Further, even with the best practices for deploying tags [25], the relative orientation between tag and animal can change during a deployment (i.e. tag slide on the animal, we refer to this event as a tag **shift**). Identifying these shifts in orientation is essential for determining the correct relative orientation. Currently, relative orientation tends to be determined manually or heuristically using portions of data where the animal’s orientation can be inferred. For example, when an animal breathes at the surface (surfacing), it is assumed that the pose should not have significant roll. A surfacing event can be inferred from pressure data and the roll estimate can then be corrected accordingly [1, 8, 26]. Tag orientation shifts tend to be identified by human inspection using sensor streams [1, 26], like accelerometer data, to identify features, such as an impact to the tag by another animal or object. However, the tag may be impacted by hydrodynamic forces, which can result in slowly changing orientation (slides) that are not captured by an accelerometer. Simulation and experimental studies have been used to estimate hydrodynamic forces that are acting on tags or imparted to the animal, but it is difficult to predict when the combined hydrodynamic and inertial forces resulting from animal motion will result in a slide [27–29]. Currently, an automated approach to identify these relative changes in orientation is lacking.

Toward addressing these gaps, this paper presents an automated data processing pipeline (and software) that takes advantage of common characteristics of pose and gait of cetaceans to (1) Identify time instances associated with the occurrences of a relative orientation change between the tag and animal; (2) Identify the relative orientation of tag wrt the animal’s body for the identified data segment; (3) Extract gait parameters that are invariant to animal pose. The authors used biologging tag data from bottlenose dolphins, humpback whales, and beluga whales to validate and demonstrate the methods. The proposed analysis approach will facilitate the use of biologging tags to study cetacean locomotion and behavior, with the desire to augment the community’s knowledge base of data processing methods and software. The method can be used directly with cetacean data from any tag platform equipped with an accelerometer, magnetometer, and pressure sensor. Discussion and suggestions related to data processing best practices are also provided. The MATLAB source code and presented data are publicly available (https://github.com/ding-z/cetacean-pose-gait-analysis.git).

### The Demonstrating Biologging Tag Platforms: MTag and DTag

The main biologging tag considered in this work is the MTag platform (Figure 1-Left) designed and built by the ESTAR Lab led by Dr. K. Alex Shorter at the University of Michigan. The suction cup-based tag has electronics encapsulated in 3D printed housings. Sensors onboard include a 9 DOF (Degrees of Freedom) IMU (Inertial Measurement Unit) with an accelerometer, gyroscope, magnetometer, and additional sensors to record temperature and pressure. The board also supports the addition of an external 1-DOF Hall effect sensor, which functions as a flow speed sensor when used in conjunction with a magnetic micro-turbine mounted outside the tag housing [30]. The tags were configured to record the IMU at 50 Hz and all other sensors at 10 Hz. To be minimally invasive and to minimize deployment complexity, the tags are attached to the animals with a set of four silicone rubber suction cups. As a result, deployments can be done within a few seconds.

Methods presented in this paper also support other cetacean tag platforms equipped with an accelerometer, magnetometer, and pressure sensor. For example, the DTag platform [1] has been commonly used for wild cetacean deployments (Figure 1-Right). The DTag also attaches to an animal via suction cups and contains an accelerometer, magnetometer, pressure sensor, high-performance hydrophone, but no gyroscope. The presented methods are demonstrated with MTag data from bottlenose dolphins and DTag data from humpback whales and beluga whales.

### Pose-Gait Patterns in Tag Data and Assumptions

The proposed work uses biologging tag data (from a 3-axis accelerometer, 3-axis magnetometer, and pressure/depth sensor) and the general data patterns associated with a cetacean gait to estimate animal pose. While an accelerometer measures acceleration due to both gravity and motion, we apply a simplified approach in this work and use the low-pass filtered 3-axis accelerometer data as the measurement associated with gravity in the tag coordinate frame (***A***^(*tag*)^). The magnitude and direction of the gravity force (vertically downwards) are known, and the resulting tag measurement always points vertically upwards in the world coordinates.

As the animal changes pose, the amount of the gravitational force measured by each of the component axes of the accelerometer changes in the tag coordinate frame (***A***^(*tag*)^), but the magnitude of the total signal is constant. We use ***A***^(*tag*)^, together with the rate of the depth change, to reveal the pattern associated with the animal’s gait, and we refer to the plot as the **orientation sphere** (Figure 2, to be distinguished from the “orientation sphere” presented in [31] for animal head movement visualization). Even though the animal could be in any pose at a particular time, common characteristics are assumed about the general gait patterns.

**Fig 2.**
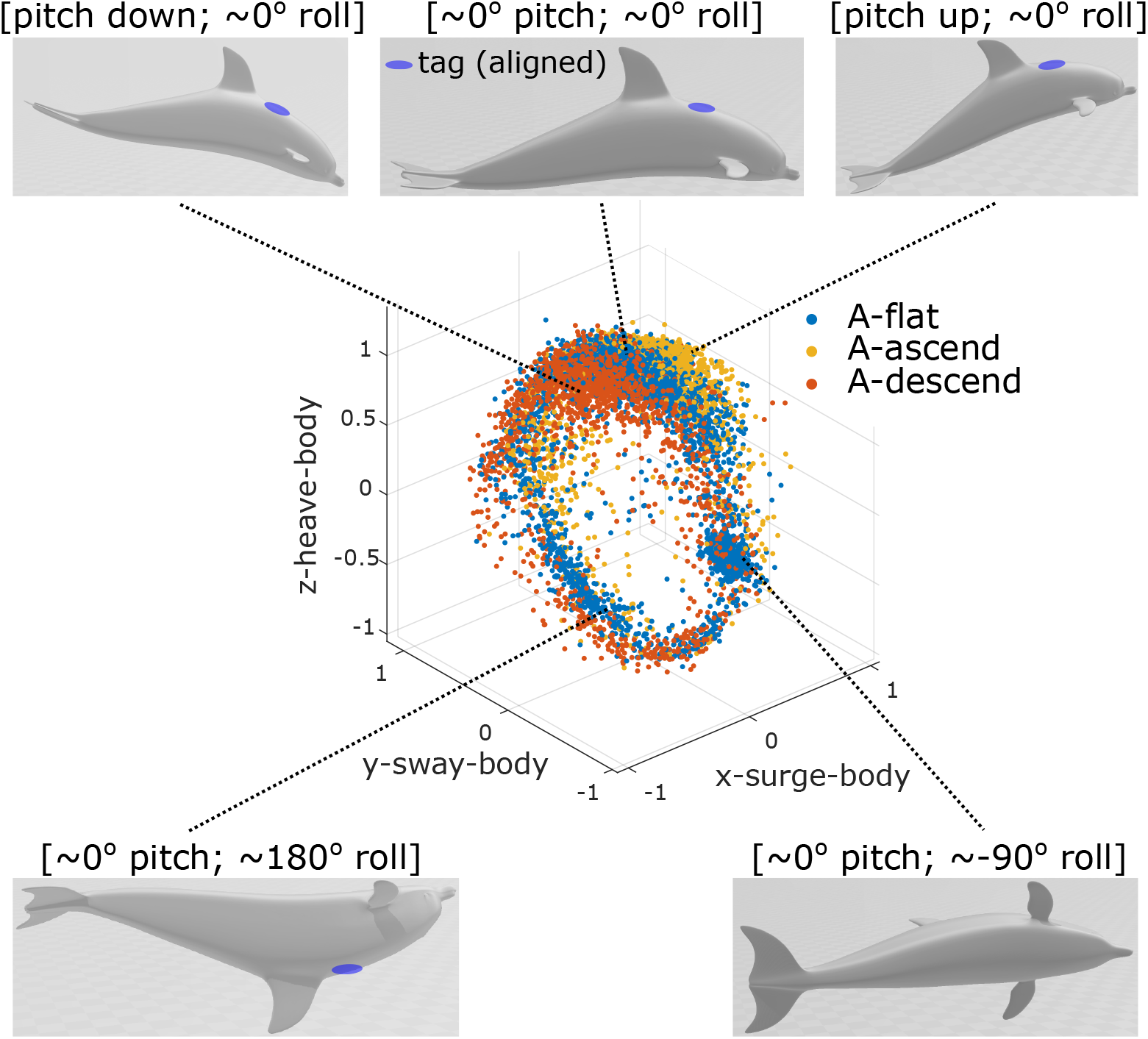
An **orientation sphere** for a bottlenose dolphin with the biologging tag aligned with the animal’s body, (i.e., data in the animal’s body coordinates without the need for a correction). Each data point represents the acceleration measurement at one time instance, and is clustered into ‘flat,’ ‘ascend,’ or ‘descend’ depending on its depth speed. The acceleration measurement is dominated by gravity and appears to point vertically up in the world coordinates constantly but changes direction in the animal’s body coordinates depending on the pose of the animal at the time. The swimming motion of an animal is composed of a sequence of poses, which corresponds to particular areas of the orientation sphere (e.g., the top 3 poses cover a shallow diving cycle and correspond to the top area of the orientation sphere).

In particular, three assumptions are made about the gait of the animal and the associated data pattern under *general* situations. The first and most important assumption is that cetaceans swim with a neutral roll angle more often than a leftward or a rightward roll. Second, it is assumed that the animal has a positive pitch when ascending and a negative pitch when descending. Third, the pattern of one signal segment is assumed to match another segment for the same animal. In this work, tag shift detection and orientation correction are based on these assumptions. More discussion shall be made about the solidness and influence of these assumptions through the rest of the paper.

### Nonparametric Tag Shift Detection

The third assumption stated above, that the pattern of one signal segment matches another segment for the same animal, will be violated when the relative orientation of the tag to the animal changes, i.e., the tag **shifts**, Figure 3. Even though significant animal gait changes could also violate the assumption, tag shift detection is achieved by identifying the time instances when signal patterns change. At a higher level, the method focuses on comparing data distributions from different data segments such that ‘abnormal’ segments in the dataset can be located. The comparisons are performed on individual data points without prior/expert knowledge of the specific data distribution.

**Fig 3.**
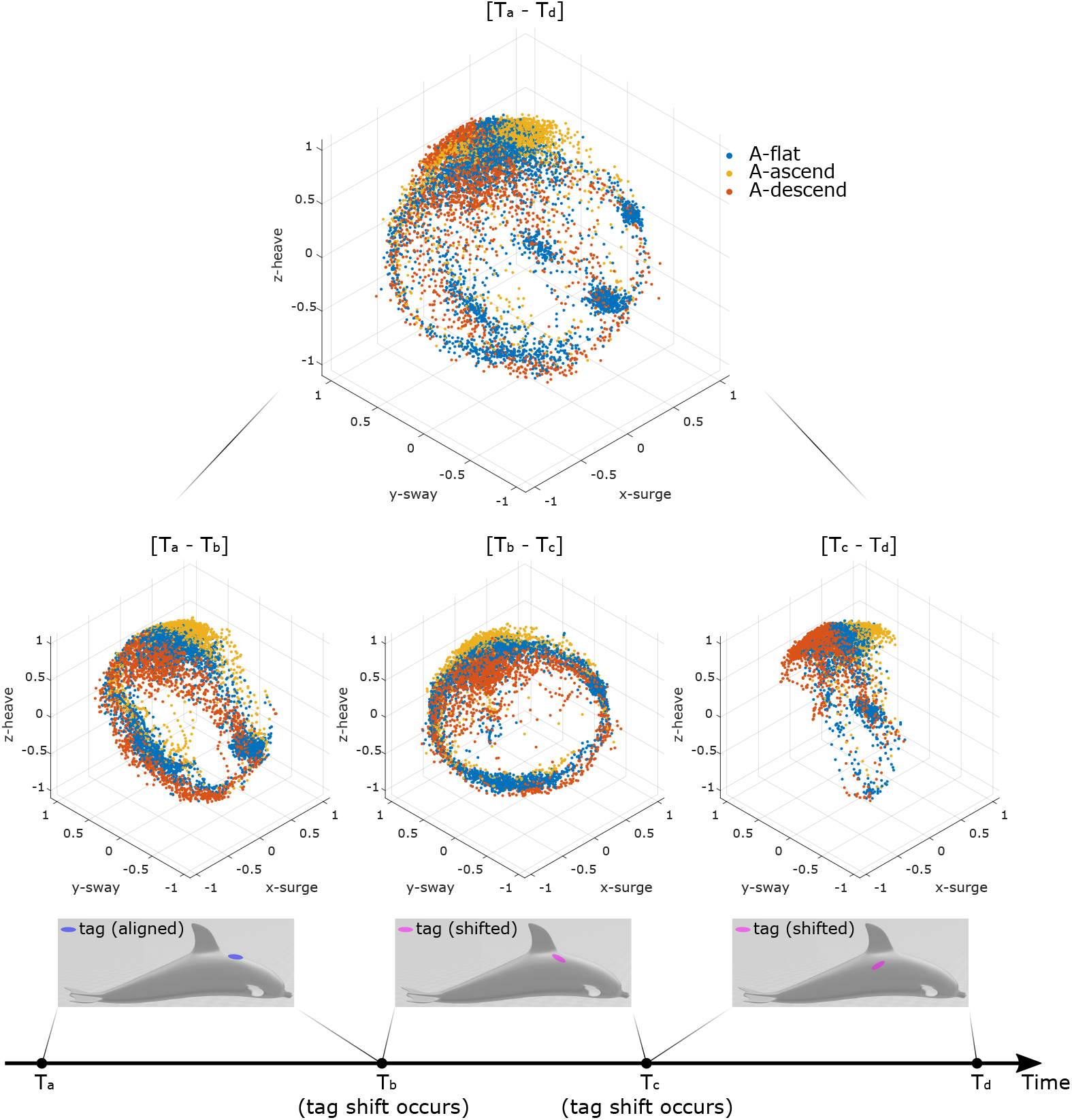
**Orientation spheres** for a bottlenose dolphin over different time horizons with the data being in the tag coordinates. Tag shifted twice in the demonstrated example with the shift time instances being *T*_*b*_ and *T*_*c*_ respectively. It is desired to 1) detect and identify the shift instances *T*_*b*_ and *T*_*c*_, and 2) transform the data from the tag coordinates to the animal’s body coordinates for all data segments (i.e. [*T*_*a*_-*T*_*b*_], [*T*_*b*_-*T*_*c*_], and [*T*_*c*_-*T*_*d*_]).

Before solving the entire shift detection problem, we first describe our approach to a more constrained subproblem: how to identify a tag shift, if it exist, within a given signal segment *S* that contains at most one shift. For this subproblem, we equally divide *S* into temporally adjacent segments *S*_1_ and *S*_2_, each has duration *D*_*s*_ (e.g. *D*_*s*_ = 10 minutes), knowing that a shift could lie in either *S*_1_ or *S*_2_, but not both. Without loss of generality, we assume the shift lies in *S*_2_. *S*_1_ then serves as the template for identifying the time instance *t*_2_ when the shift happens in *S*_2_. With *t*_2_, *S*_2_ can be further divided into 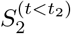 and 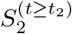. Then, we know 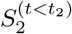 shares the same distribution as *S*_1_ and 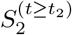 diverges from *S*_1_.

The question now is how to locate such *t*_2_ in *S*_2_, using *S*_1_ as the template. For each data point in *S*_2_, we find its *K* (see footnote^1^) nearest spacial neighbors in *S*_1_ and compute the average distance from the point to its neighbors. If this distance is within an empirically defined threshold, the data point in *S*_2_ is considered an **inlier** of *S*_1_; otherwise, it is classified as an **outlier**. After this process, all of the points in *S*_2_ are assigned with a value of 1 (inlier) or 0 (outlier). A temporal moving average filter is applied to these 1s and 0s to obtain a local inlier percentage (*InPct*) value. As illustrated in Figure 4, the time instance *t*_2_ is found by identifying the first time *InPct* drops below a defined threshold. If *InPct* never drops below or *t*_2_ is close to the boundary between *S*_1_ and *S*_2_, this indicates that a shift could be lying in *S*_1_ instead of *S*_2_. This procedure can then be applied to check *S*_1_ by using *S*_2_ as a template, aiming at finding time instance *t*_1_ in *S*_1_ that corresponds to a shift.

**Fig 4.**
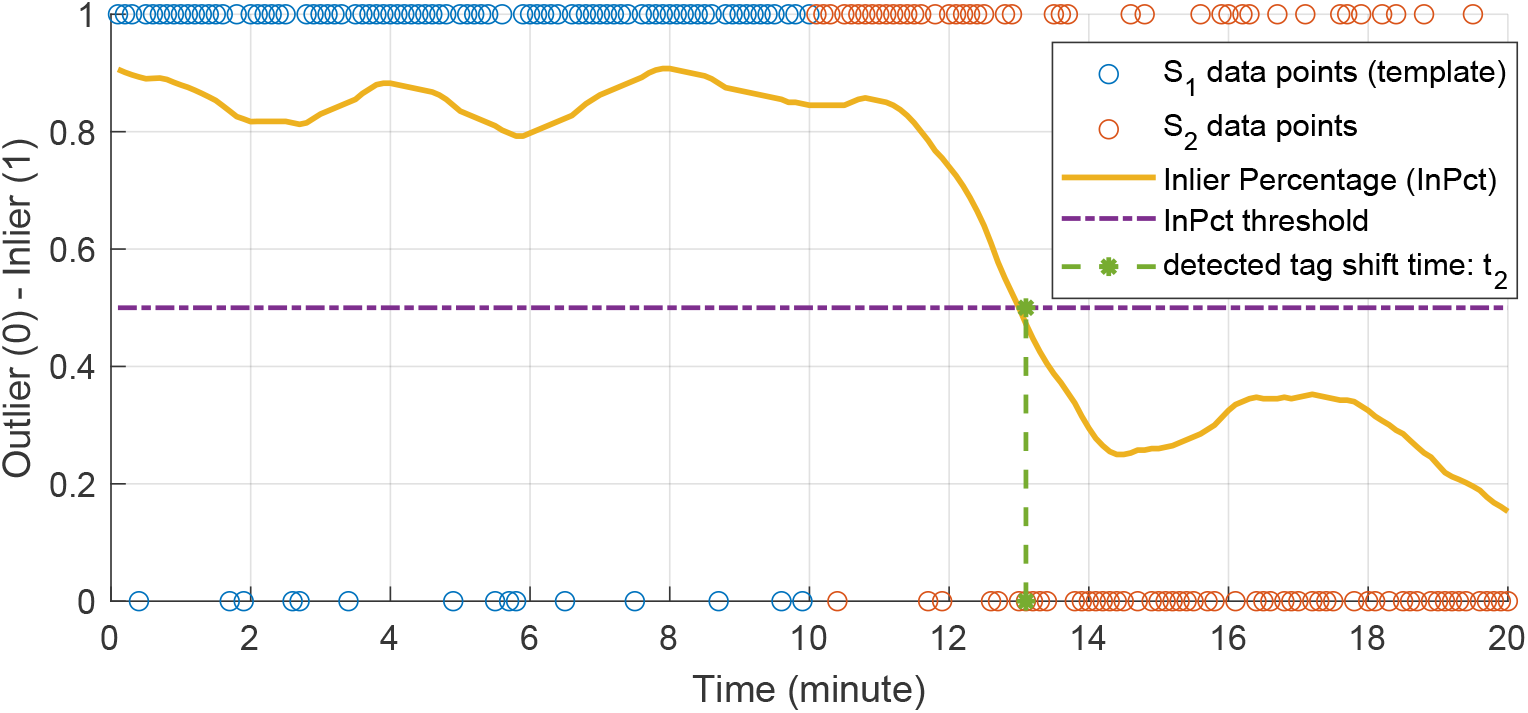
Conceptual illustration of finding tag shift time *t*_2_ in data segment *S*_2_ using data segment *S*_1_ as a template. Both segments share the same duration *D*_*s*_ = 10 minutes. Each data point from either *S*_1_ or *S*_2_ can be checked against the template distribution (*S*_1_) to decide whether it is an inlier (1) or an outlier (0) of the template. An inlier percentage (*InPct*) value can be calculated over time, and the tag shift time can be located by finding when *InPct* drops below an empirically defined threshold. If *t*_2_ does not exist or it is close to the boundary between *S*_1_ and *S*_2_, another step would be taken to use *S*_2_ as the template to check for a potential shift time *t*_1_ in *S*_1_.

With a strategy defined above to find one (or zero) shifting instance within two segments, *S*_1_ and *S*_2_, an algorithm for finding all tag shifts is presented in Figure 5. Based on the procedures described above, the algorithm starts with adjacent data segments *S*_1_ and *S*_2_. Both segments share a fixed empirically defined duration *D*_*s*_ that reflects the expected tag shift interval (e.g. *D*_*s*_ = 10 minutes). If neither segment contains a shift (branch 10), then *S*_1_ remains unchanged, and *S*_2_ moves forward to the following data segment to check for a shift. Once a shift is detected, both segments will be redefined such that *S*_1_ starts from the identified shift and *S*_2_ adjacently follows (branches 6 to 9).

**Fig 5.**
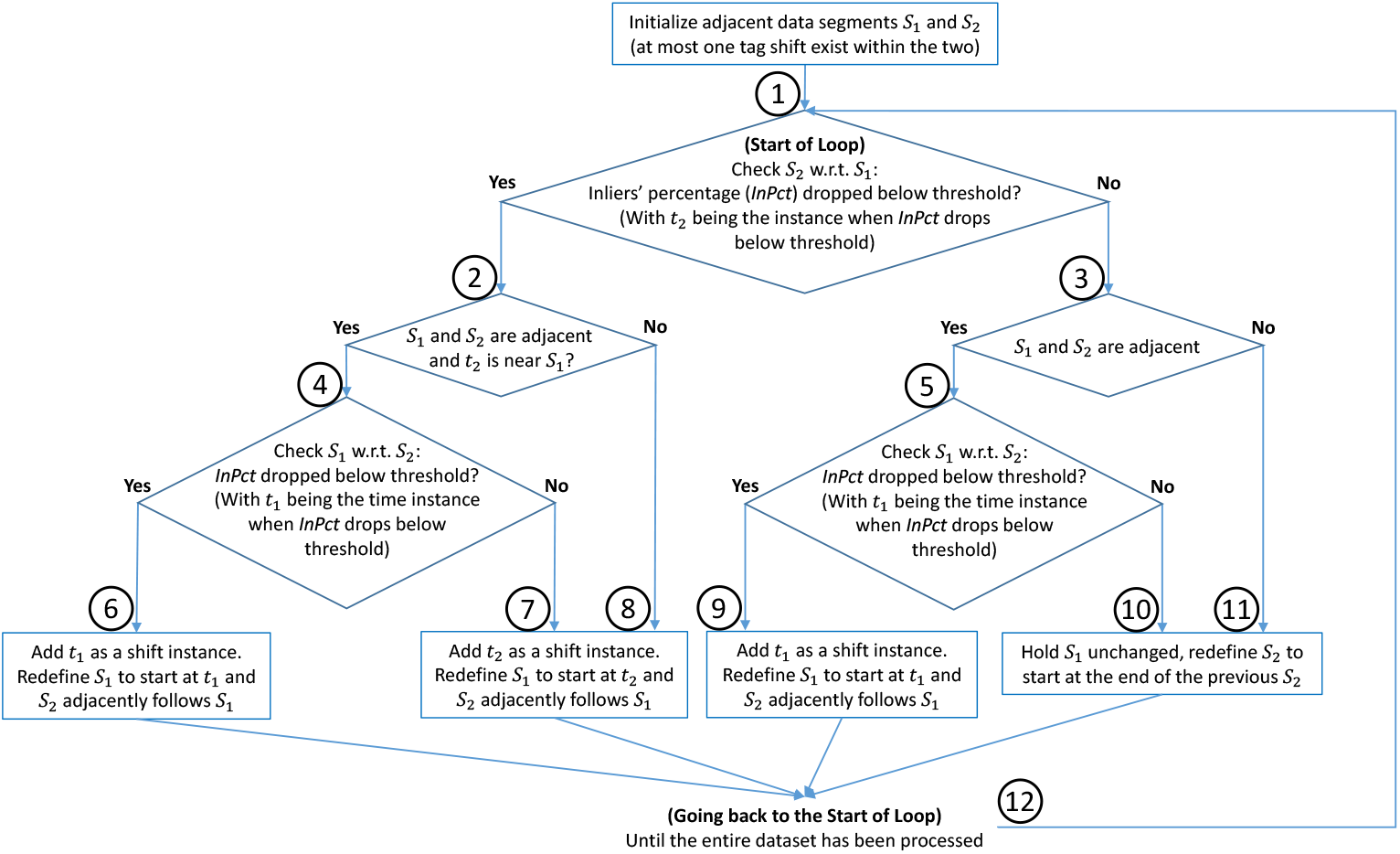
Tag shift detection algorithm. Each branch is marked by a circled number (1 to 12).

Every time *S*_1_ is redefined (i.e. *S*_1_ and *S*_2_ are adjacent), the algorithm searches for a shift to exist in either *S*_1_ or *S*_2_, starting with *S*_2_ (branch 1). If *S*_2_ does not contain a shift (branch 3), then the algorithm checks *S*_1_ for a shift (branch 5, followed by branches 9 & 10). If *S*_2_ contains a shift instance *t*_2_ (branch 2), and *t*_2_ is not near (see footnote^2^) *S*_1_ (branch 8), then *t*_2_ is recorded as a shift. However, if *t*_2_ is near *S*_1_ (branch 4), the algorithm will also check *S*_1_ to decide whether the shift is lying in *S*_1_ but appears to be *t*_2_ initially. Then either *t*_1_ or *t*_2_ will be recorded as the shift, depending on the search result of *S*_1_ (branches 6 & 7). When only *S*_2_ is redefined (i.e. *S*_1_ and *S*_2_ are not adjacent), the algorithm only searches for a shift in *S*_2_ (branches 8 & 11). These steps are repeated until the entire dataset has been searched (branch 12).

### Pattern-Based Orientation Correction

For a given segment of data that contains no tag shift but has an unknown tag-animal configuration, the tag data from the accelerometer and magnetometer (and gyroscope, if equipped) can be rotated (i.e., change of coordinates) to transform the data from tag coordinates to the animal’s body coordinates. In this work, such a rotation is found by matching the measured general motion pattern, i.e., orientation sphere, to the assumed one, as discussed in this section. Depth measurement is used to group the tag coordinates’ gravity measurements ***A***^(*tag*)^ into three clusters: 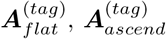 and 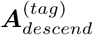 in accordance with the animal swimming horizontally, ascending, and descending.

We further define the **dominant direction** as the gravity measurement direction associated with the most common pose of the animal for each of the three swimming conditions: 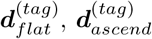, and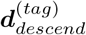. By the assumptions about the general motion of these animals (Section *Pose-Gait Patterns in Tag Data and Assumptions*), the most common pose of a cetacean under each of the three swimming conditions is assumed to be known. In particular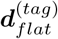 would correspond to the gravity measurement when the animal has zero pitch and zero roll, while 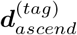 corresponds to positive pitch and zero mroll, and 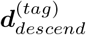 corresponds to negative pitch and zero roll. In other words, we know how the dominant directions are orientated in the animal’s body coordinates.

Specifically, 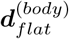 is aligned with the body’s *z*-axis (*z*^(*body*)^), and 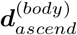 lies in the first quadrant of the plane formed by the *x*^(*body*)^ and *z*^(*body*)^ axes, while 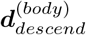 lies in the second quadrant (Figure 6-Top Row). So if 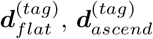 and 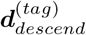 can be found in the tag coordinates (Figure 6-Bottom Row), the rotations that map them to 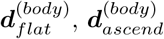 and 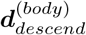 can be found and applied to map other data from tag coordinates to animal body coordinates (Figure 6).

**Fig 6.**
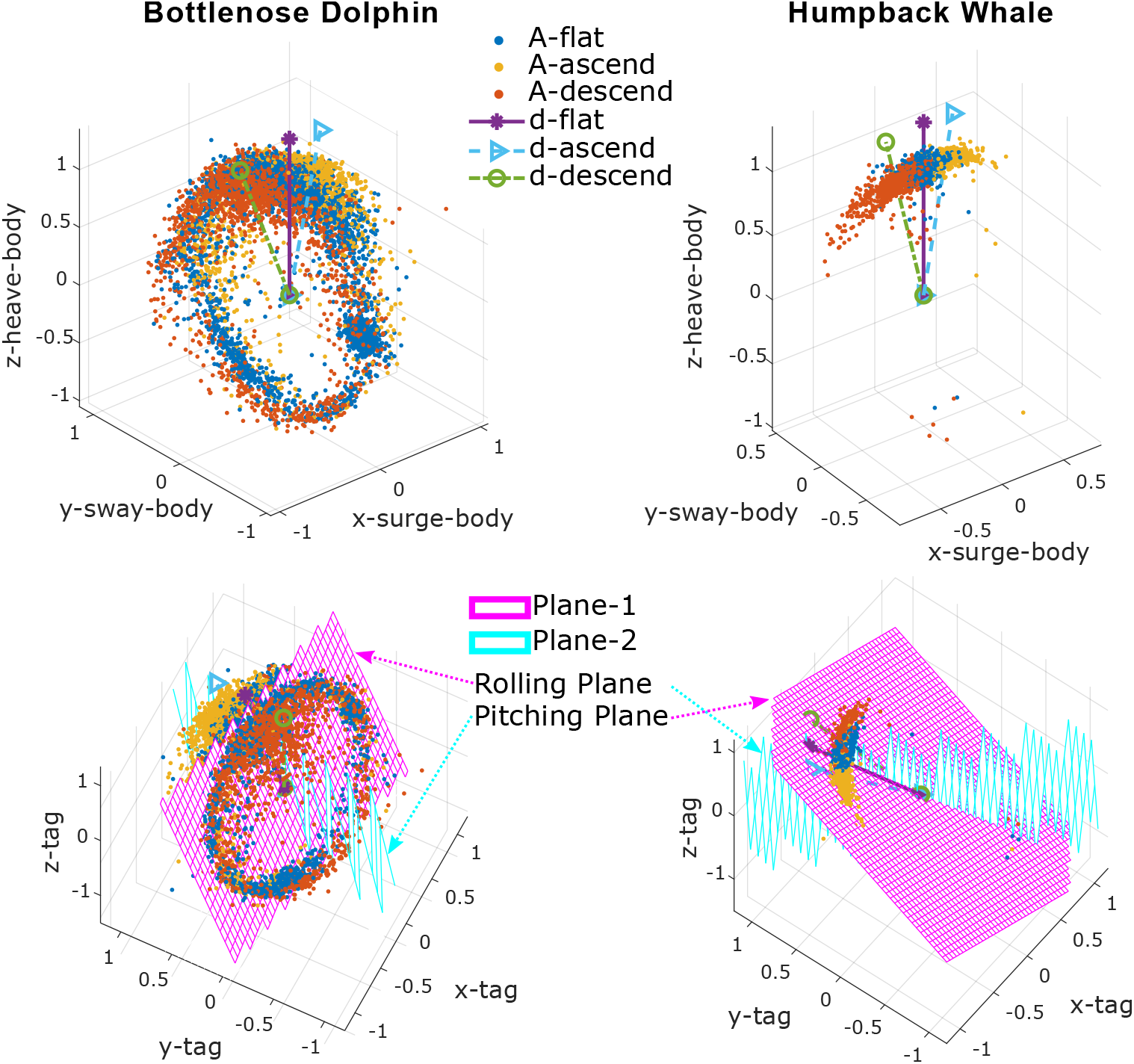
**Orientation spheres** for a data segment of a bottlenose dolphin (**left column**) and a humpback whale (**right column**). The plots visualize the orientation correction method applied to an uncorrected data segment to find the dominant directions in the tag’s coordinates (**bottom row**). The prevailing directions can then be mapped to their assumed directions in the body’s coordinates (**top row**).

To determine the dominant directions within uncorrected data, two perpendicular planes are fitted to ***A***^(*tag*)^ using the RANSAC algorithm [32] (Figure 6-Bottom Row), which is particularly robust to outliers and imbalanced data, as compared to least square based approaches. The two planes essentially correspond to two physical motions: rolling with zero pitch for the **rolling plane** and pitching with zero roll for the **pitching plane**.

Firstly, plane-1 (through the origin) is fitted to the most significant distribution in the data, which could be either 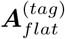 (e.g. Figure 6-Bottom Left, plane-1 is a **rolling plane**) or 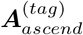 and 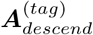 (e.g. Figure 6-Bottom Right, plane-1 is a **pitching plane**), depending on the animal and environment. Secondly, plane-2 (through the origin) is fitted to the rest of the data under the constraint that it is perpendicular to plane-1. The toolbox can assign the roles of **rolling plane** and **pitching plane** to the two planes automatically.

The intersection line of the two planes is colinear with 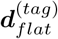 while both 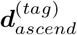 and 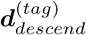 lie in the **pitching plane**. If we further define 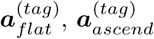 and 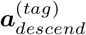 to be the average values of 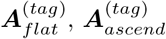, and 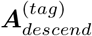, then projecting 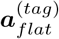 onto the intersection line produces 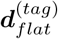, while projecting 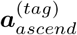 and 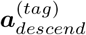 onto the **pitching plane** returns 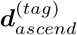 and 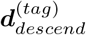(Figure 6-Bottom Row), respectively.

With 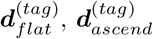, and 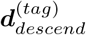 identified in the tag coordinates, the data can be put into the animal’s body coordinates (i.e. ***A***^(*tag*)^ → ***A***^(*body*)^) with two rotations: first, a rotation (***R***_1_) to align 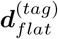 with the body’s *z*-axis (*z*^(*body*)^) and second, a rotation (***R***_2_) around the *z*-axis so ***d***_*ascend*_ and ***d***_*descend*_ are on the *x*-*z* plane with ***d***_*ascend*_ on the positive *x*-axis side and ***d***_*descend*_ on the negative *x*-axis side (Figure 6-Top Row). Other data are mapped from tag coordinates to body coordinates using ***R***_1_ and ***R***_2_:

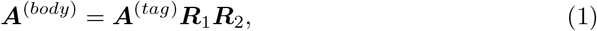

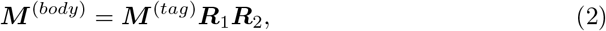

with ***A***^(*body*)^ and ***M*** ^(*body*)^ being the transformed (animal’s body coordinates) data from the accelerometer and magnetometer.

### Animal Pose Calculation and Gait Characterization

Roll, pitch, and yaw are widely used to describe the pose of animals [1, 8]. In this work we are computing them in a conventional [1] way using the transformed (animal’s body coordinates) data from accelerometer ***A***^(*body*)^ and magnetometer ***M*** ^(*body*)^. Define ***R***_*x*_(*α*) to be the rotation matrix for rotating the data about the coordinates’ *x*-axis with angle *α*. And similarly ***R***_*y*_(*β*) about the *y*-axis with angle *β* and ***R***_*z*_(*γ*) about the *z*-axis with angle *γ*. For time instance *t*, the 3-axes body coordinates’ accelerometer and magnetometer data are represented as 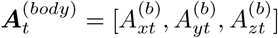 and 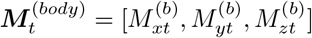. Roll, pitch, and yaw are calculated as:

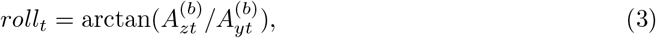

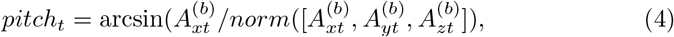

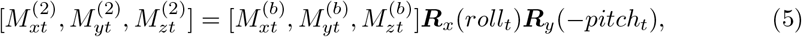

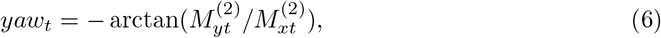

where we note that the body coordinates’ *x*^(*body*)^, *y*^(*body*)^, and *z*^(*body*)^ axes are defined to be pointing forward, leftward, and upward wrt the animal’s body, respectively (Figure 1).

Let a set of moving coordinates {*x*^(*move*)^, *y*^(*move*)^, *z*^(*move*)^} be initially aligned with the earth’s inertial coordinates {*x*^(*inertial*)^, *y*^(*inertial*)^, *z*^(*inertial*)^} (see footnote^3^). Then yaw, pitch, and roll rotate the moving coordinates from the inertial coordinates to the animal’s body coordinates in 3 steps:

1. Yaw corresponds to a positive rotation around the moving coordinates’ current *z*^(*move*)^ axis (same as the inertial *z*^(*inertial*)^ axis at the moment), rotates the moving coordinates so *x*^(*move*)^ is aligned with the projection of the animal’s *x*^(*body*)^ axis on the earth horizontal plane.
2. Pitch measures the angle between the body coordinates’ *x*^(*body*)^ axis and the horizontal earth plane, rotates around the moving coordinates’ current *y*^(*move*)^ axis and makes *x*^(*move*)^ tilted from horizontal to be aligned with the animal’s *x*^(*body*)^ axis. Note that a positive pitch (i.e., animal head up) corresponds to a negative rotation around *y*^(*move*)^, with *y*^(*body*)^ defined pointing to the left of the animal.
3. Roll represents a positive rotation around moving coordinates’ current *x*^(*move*)^ axis (which is now aligned with the animal’s *x*^(*body*)^ axis) that brings moving coordinates’ *y*^(*move*)^ axis from horizontal to the animal’s *y*^(*body*)^ axis. After this step, *x*^(*move*)^, *y*^(*move*)^, and *z*^(*move*)^ are aligned with *x*^(*body*)^, *y*^(*body*)^, and *z*^(*body*)^, respectively.

This pitch estimation is commonly used to describe the fluking gait of the animal [4, 5, 8]. But because pitch only measures the angle between the animal’s *x*^(*body*)^ axis and the horizontal earth plane, it does not capture the gait characteristics well when the animal is rolling. The situation degrades more when the animal is fluking with a roll angle of 90^*°*^ where the fluking motion barely shows up in the pitch measurement. Ideally, we want an angle measurement wrt the animal’s own *y*^(*body*)^ axis to describe its gait. Unfortunately, even though this can be done by integrating the gyroscope measured angular rate about the *y*^(*body*)^ axis of the animal, a gyroscope is not available in all tag platforms and the integration is subject to measurement drift due to accumulated errors.

Here,{*x*^(*body*)^, *y*^(*body*)^, *z*^(*body*)^} are moving wrt the earth inertial coordinates following the animal’s motion (both **high** frequency, e.g. fluking; and **low** frequency, e.g. turning from ascending to descending). We now define a new set of coordinates 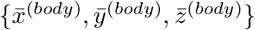 that only follow the **low** frequency motion of the animal, which effectively represents the neutral pose of the animal during high frequency periodic motion. Then 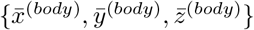 can be used to characterize animal’s **high** frequency motion (e.g. fluking) wrt the animal itself, rather than wrt the earth’s inertial coordinates.

Specifically, representing *x*^(*body*)^ in 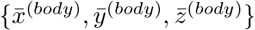 returns the dynamic heading of the animal wrt its own neutral body pose. For example, an animal’s head tilting up and down wrt its neutral body pose would result in the vector *x*^(*body*)^ swinging in the 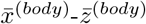 plane. We now refer to the vector *x*^(*body*)^, represented in 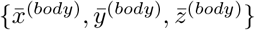, as the **vector of dynamic pose** (***V***_*dp*_):

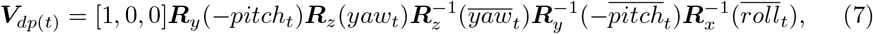

where [1, 0, 0] is an unit vector pointing forward while 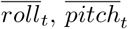 and 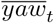 are the low-pass filtered roll, pitch and yaw values at time *t*. We note that a positive animal pitch is a negative rotation around the *y*-axis under the current axes definition. Within the equation, ‘[1, 0, 0]***R***_*y*_(−*pitch*_*t*_) ***R***_*z*_(*yaw*_*t*_)’ finds the animal’s current 3D heading in the inertial coordinates (i.e. *x*^(*body*)^ in the earth inertial coordinates). ‘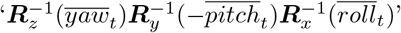 ‘ then maps *x*^(*body*)^ from the inertial coordinates to the low frequency body coordinates 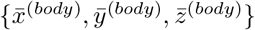.

The resulting ***V***_*dp*_ gives the dynamic 3D heading of the animal wrt the animal’s current neutral body pose (i.e. the low frequency body pose). Since the fluking motion of cetaceans is happening in the 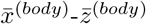 plane (i.e. **pitching plane**), the resulting ***V***_*dp*_ would also be swinging in the 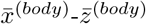 plane, regardless of the actual orientation of the animal (see footnote^4^). We further define **dynamic pitch** (*pitch*_*dp*_) and **dynamic yaw** (*yaw*_*dp*_) to be the angles between ***V***_*dp*_ and its *x*-*y* plane as well as its *x*-*z* plane, respectively:

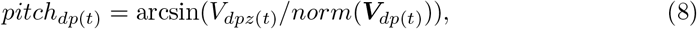

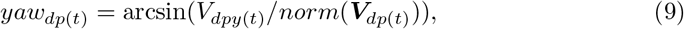

with ***V***_*dp*(*t*)_ = [*V*_*dpx*(*t*)_, *V*_*dpy*(*t*)_, *V*_*dpz*(*t*)_] at time *t*. The **dynamic pitch** (*pitch*_*dp*_) and **dynamic yaw** (*yaw*_*dp*_) approximate the animal’s dynamic angle changes around its 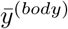 and 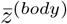 axes, respectively. The *pitch*_*dp*_ measure for a cetacean is what we desired earlier: ‘an angle measurement wrt the animal’s own body *y*^(*body*)^ axis to describe its gait.’

An animal’s per-stroke fluking period and amplitude are calculated by automatically locating and processing the positive and negative peaks in *pitch*_*dp*_, which enables a wide variety of further analyses (e.g., investigating the animal’s fluking frequency and amplitude when diving speed is greater than 2 m/s). Meanwhile, data segments that do not contain any peak are marked as passive behaviors (e.g., gliding).

## Validation

The validation objectives were to assess: (**1**) how precise can the proposed method be in identifying the tag shift instances; (**2**) how well can the proposed method correct the data to compensate for the misalignment between tag and animal; (**3**) how accurate is the calculated animal pose compared to ground truth; (**4**) how sensitive is the method in response to a small amount of tag shift; (**5**) what is the impact of the defined segment duration *D*_*s*_ to the shift detection performance; and (**6**) how applicable is the technique for different cetaceans.

Biologging tag (MTag) data collected from bottlenose dolphins (*Tursiops truncatus*) under human care in Dolphin Quest Oahu Hawaii were used for quantitative validation of the methods. The biologging tag was aligned with the animal and attached 20 cm behind the blowhole non-invasively via four silicone suction cups (Figure 1-Left).

As a correctly aligned tag measurement differs from a misaligned tag primarily by a rotation to the data (i.e., a change of coordinates), we injected artificial rotations to the initially aligned data to simulate the effects of tag shifts. For validation purposes, randomly designed rotations were applied to each randomly defined data segment to simulate the effects of a tag shift. More specifically, 18 datasets, with an average duration of 87 (± 23) minutes, were included for the validation, and 100 random simulations were conducted for each dataset. The dataset was randomly broken into *k* + 1 segments for each simulation run, with the random integer *k* ranging between 1 and 6, representing the number of injected tag shifts. A random rotation, with both random direction and magnitude (in degree), was then applied to each data segment to simulate the effect of a tag shift. The method uses an empirically specified segment duration setting *D*_*s*_ = 10 minutes for shift detection.

For tag shift detection (**Objective 1**) we define **error** of a detection as the absolute time difference between the detected shift instance and the nearest injected shift instance. Further, the detection is a **positive detection** if the error is within 300 seconds (5 minutes), otherwise a **negative detection**. Then **precision** and **recall** are calculated as:

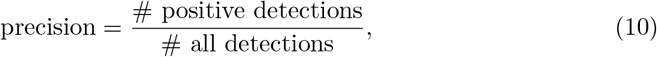

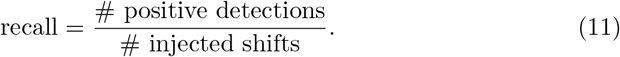

To assess whether the random rotations have been corrected well (**Objective 2**), poses (roll, pitch, and yaw) calculated from **corrected** simulated data segments (condition **A**) were compared with poses calculated from the original data segments (no random rotation injected, condition **B**).

Meanwhile, to evaluate the stated pose calculation method (**Objective 3**), the poses calculated using the stated method (involved accelerometer and magnetometer, condition **B**) were further compared with the poses calculated using a gradient descent based filtering approach [24] (referred to as the *Madgwick* ‘s approach after the author’s name), which used accelerometer, magnetometer, and **gyroscope** data (condition **C**).

Further, to assess the sensitivity of the proposed methods (**Objective 4**), we repeated the computation experiments by injecting simulated tag shifts, with the simulated tag rotations having random directions (same as before) but **fixed** degrees for each run. Specifically, 50 random runs were made to each dataset for each fixed degree. For each run, *k* + 1 rotations were injected into the dataset (as before), and each rotation had a random direction with a fixed degree. The average precision and recall of the method in detecting the shifting instances were calculated for each fixed degree over all datasets and random runs. In addition, the absolute angle differences between poses calculated from corrected simulated data segments (condition **A**) and poses calculated from the original data segments (condition **B**) were calculated for each fixed angle.

The computation experiments were repeated one more time to assess the impact of segment duration *D*_*s*_ on the shift detection performance (**Objective 5**). Different *D*_*s*_ choices were evaluated against varying number of injected tag shifts (i.e. *k*) to explore the relationship between user specified segment duration *D*_*s*_ and the expected tag shift interval (determined by *k*). For each {*D*_*s*_, *k*} combination, 50 random runs were made over all datasets to calculate the average precision and recall of shift detection.

In addition to the bottlenose dolphin datasets, DTag [1] data from a wild humpback whale (22.03 hours) and a beluga whale (2.25 hours) were included to evaluate the method and demonstrate the gait analysis capability (**Objective 6**).

## Results

Among all 100 random runs over the 18 datasets (**Objective 1**), the average precision for shift instance detection is 0.87, the average recall is 0.89 while the average error is 37.5 (±18.4) seconds. The poses (roll, pitch, and yaw) calculated from **corrected** data segments (condition **A**) were compared with poses calculated from the original data segments (condition **B, Objective 2**), with results shown in Table 1-Left. The average errors and standard deviations were within 11 degrees for all poses. Meanwhile, the poses calculated in condition **B** were further compared with the poses calculated using the *Madgwick* ‘s approach [24] (condition **C, Objective 3**, Table 1-Right). The average errors were all within 3 degrees, and the standard deviations were within 6 degrees.

**Table 1.**
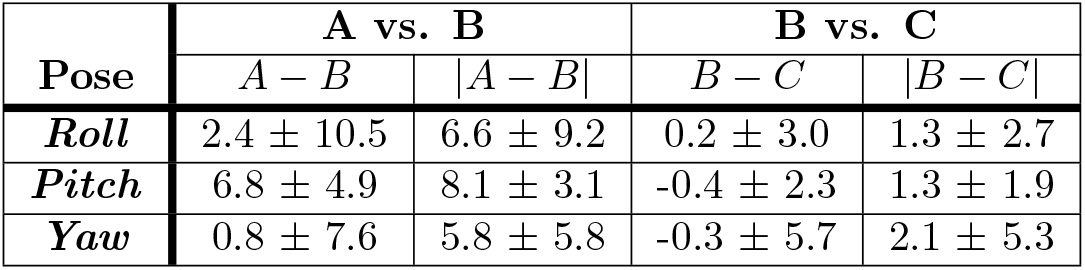
Resulting pose angle differences with different conditions (i.e. **A, B**, and **C**). Condition **A** represents pose calculated from simulated data after the simulated tag shifts have been corrected using the proposed method. Condition **B** presents pose calculated from the original data (without tag shift). Condition **C** stands for pose calculated from the original data (without tag shift) using Madgwick’s filter [24], which involves the additional use of a gyroscope. The direct difference (e.g., *A*− *B*) is used to detect a bias in the difference, while the absolute difference (e.g., | *A* − *B* |) returns the magnitude of the difference. Pose angle differences are in degree, and each cell gives a mean ± standard deviation.

To assess the sensitivity of the proposed methods (**Objective 4**), the computation experiments were repeated with virtually injected tag rotations that had random direction and **fixed** degrees. The average precision and recall for the detection of these injected tag shift instances are shown in Figure 7-Left. With a 10 degrees rotation offset, less than 30% of the injected shifts were detected, while with a 20 degrees offset, ∼54% of the shifts were detected with a detection precision of ∼79%. When the offset reaches 30∼ 40 deg, the method’s performance starts being more reliable with a recall in the 70 ∼ 80 percentile and precision being ∼85%. When the offset reaches 50 degrees and above, the method’s performance converges to ∼90% precision and recall.

**Fig 7.**
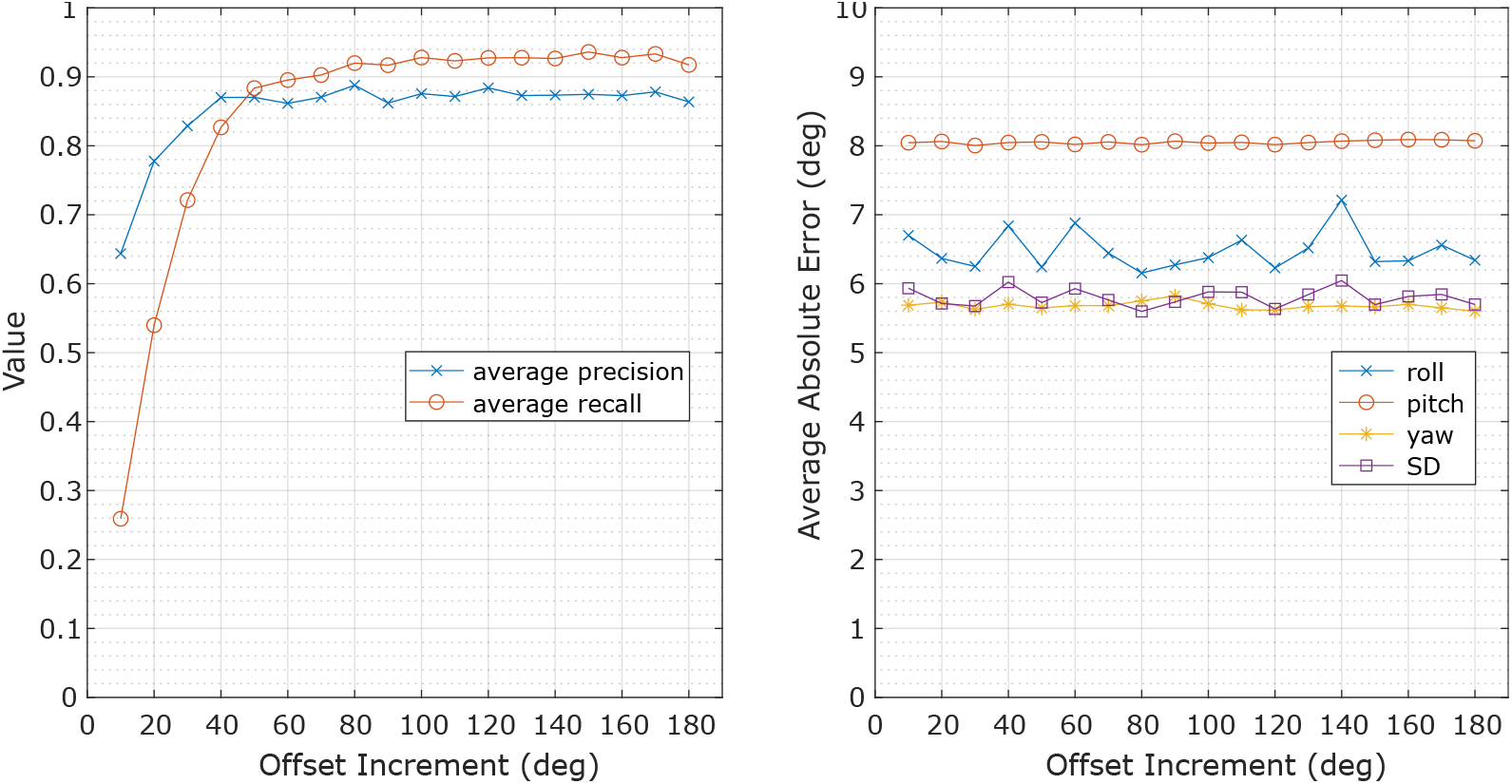
Average precision and recall of tag shift detection (**left**) and the average absolute error of the calculated animal pose after tag orientation correction (**right**) over simulated tag shifts with random direction and fixed degrees (*x*-axis). SD denotes the average of the standard deviation values for roll, pitch, and yaw at each offset increment.

Figure 7-Right presents the average absolute errors of the calculated poses after rotation correction for each of the data segments transformed by a rotation. The errors demonstrate similar values across all rotation offsets, with errors for roll being ∼6.5 degrees, pitch being ∼8.0 degrees, yaw being ∼5.7 degrees, and the average standard deviation being ∼5.8 degrees.

Figure 8 demonstrates the average precision and recall of tag shift detection using different segment duration setting *D*_*s*_ in response to varying number of injected shifts *k* (or equivalently, the average shift interval, **Objective 5**). The average precision of all *D*_*s*_ choices remain between 0.83 and 0.97, except when *D*_*s*_ = 5 minutes, whose average precision increased from 0.47 to 0.72 as the number of injected shifts increases. The average recall of *D*_*s*_ choices demonstrate a dropping trend as the number of injected shifts increases, and the drop is more significant with bigger *D*_*s*_ choice. However, the average recall values remain above ~ 0.8 when the specified *D*_*s*_ is smaller than the average shift interval.

**Fig 8.**
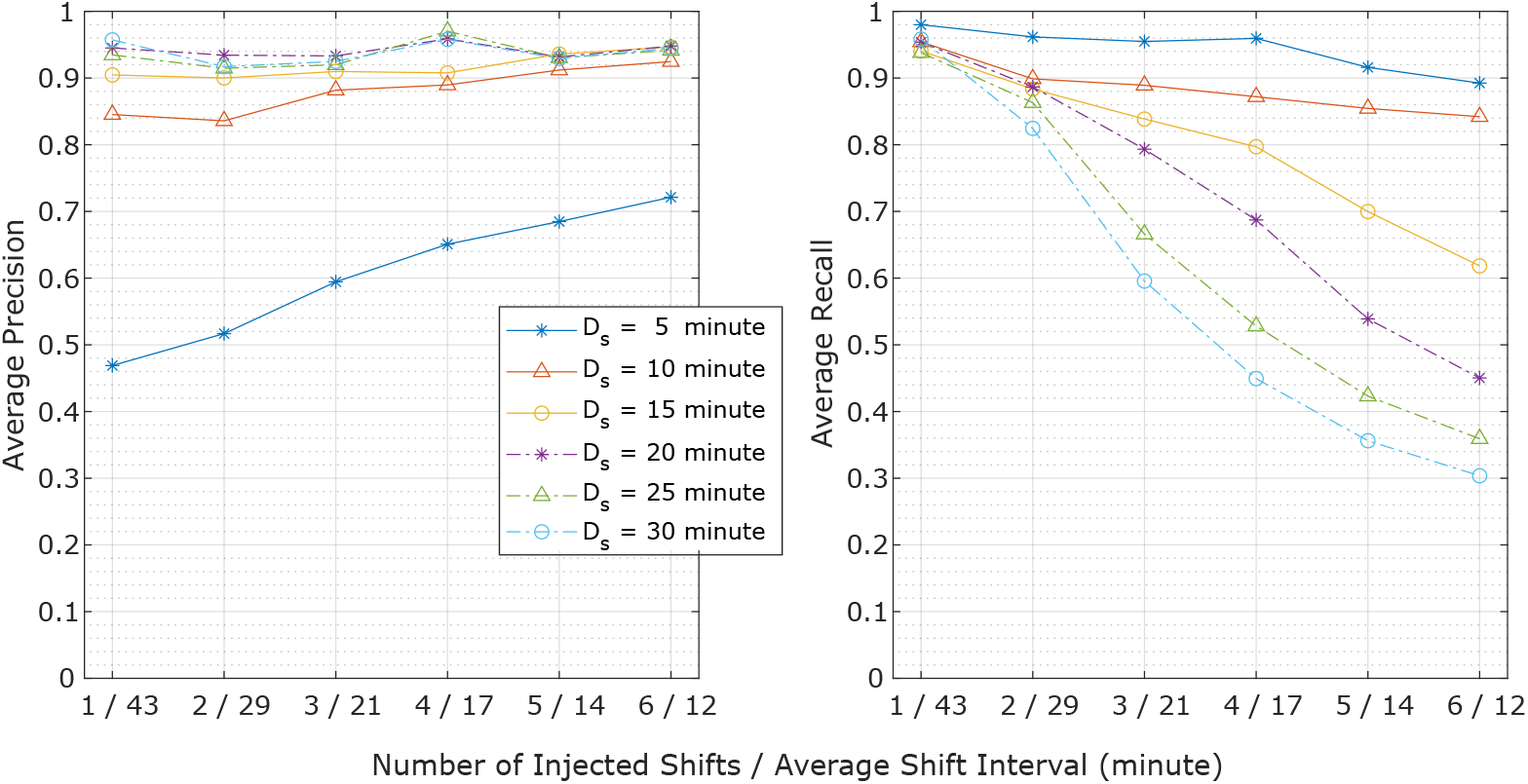
Average precision (**left**) and recall (**right**) of tag shift detection performance during simulation plotted against varying number of injected shifts (or equivalently, the average shift interval). Shift detection performance at specific segment durations (*D*_*s*_) are demonstrated by the individual curves in the plot.

Figure 9 presents the orientation spheres of an example bottlenose dolphin dataset (left column), a humpback whale dataset (center column), and a beluga whale dataset (right column, **Objective 6**). The top row shows the orientation sphere after all corrections. The middle row provides the raw data before tag shift detection and orientation corrections. And the bottom row shows the orientation method being applied to one of the data segments to identify the dominant directions. For example, the raw dataset of the bottlenose dolphin (middle row - left) contains 4 simulated tag shifts, while 6 tag shifts were detected from the wild humpback whale dataset (middle row - center). The raw tag data from the wild beluga whale (middle row - right) indicates that the tag was not aligned with the animal but contains no detected tag shift (i.e., no relative motion between tag and animal was detected).

**Fig 9.**
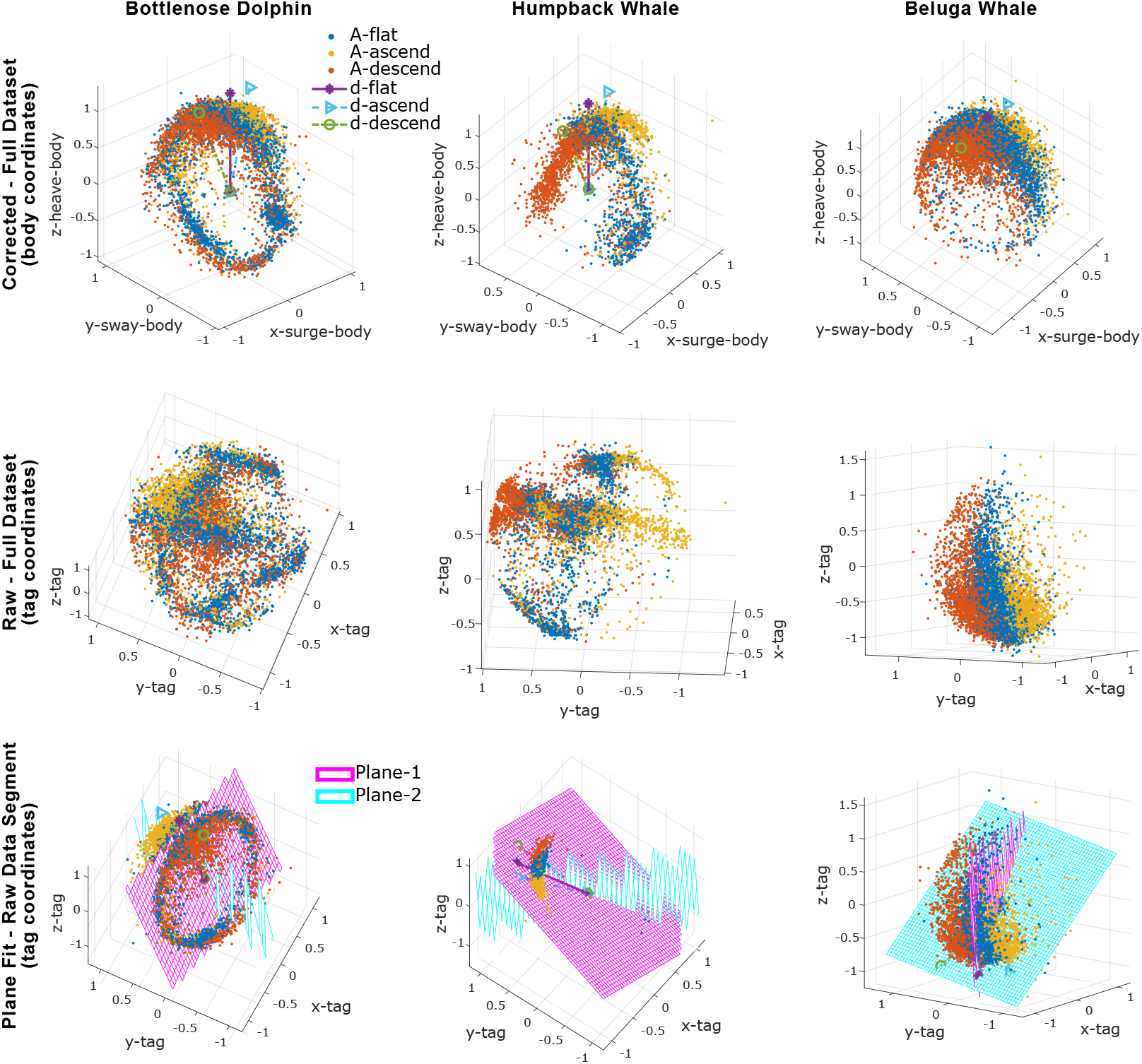
**Orientation spheres** for a bottlenose dolphin (left column), a humpback whale (center column), and a beluga whale (right column). The plot of accelerometer data clustered by depth speed is referred to as **orientation sphere**. After detecting tag shifts and correcting tag orientation misalignment, the top row of subplots presents the orientation spheres in the animal’s body coordinates. The middle row shows the data in the tag coordinates before any correction. The bottom row visualizes the orientation correction method being applied to an uncorrected data segment. The dolphin dataset (left column) contains 4 simulated tag shifts. The humpback whale dataset (center column) has 5 detected tag shifts. While the beluga dataset (right column) has no detected tag shifts (i.e., no relative motion between tag and animal was detected).

Figure 10 presents example data segments from one of the bottlenose dolphin (left) and the beluga whale (right) datasets, where the pose of the animal was calculated from the corrected body coordinates’ data and the gait of the animal was characterized using the *pitch*_*dp*_ (**dynamic pitch**) of the animal. The mean fluking period of this bottlenose dolphin is 1.0 s with an average amplitude of 14.1 degrees. The mean fluking period of the beluga whale is 1.6 s with an average amplitude of 16.8 degrees. For the humpback whale dataset, the mean fluking period is 6.7 s with an amplitude of 12.2 degrees.

**Fig 10.**
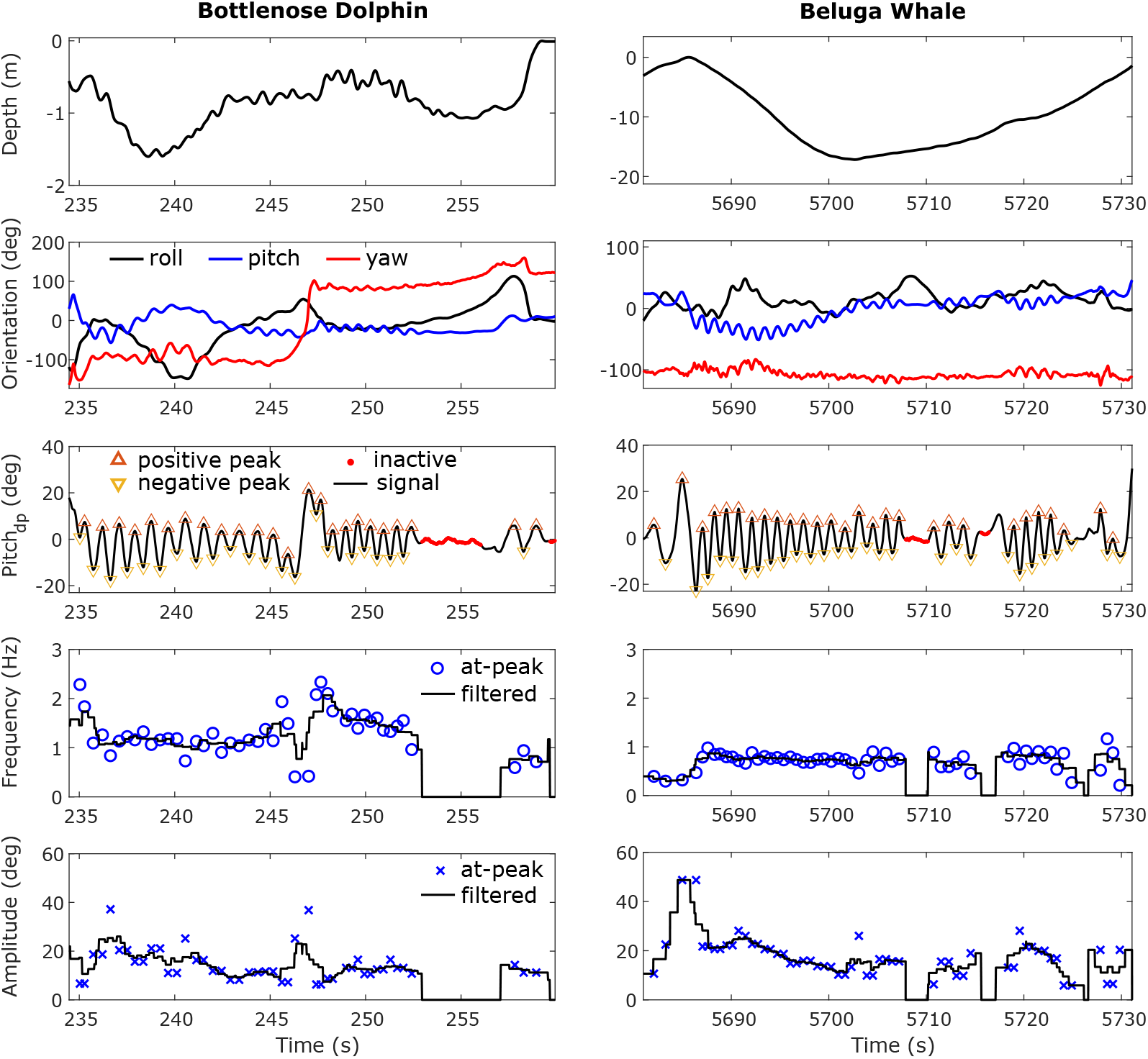
Example data segments from a bottlenose dolphin (**left**) and a beluga whale (**right**) with animal’s gait parameterized via **dynamic pitch** (*pitch*_*dp*_). When the animal has a big roll angle (e.g., during the time around 240 seconds in the bottlenose dolphin dataset), the fluking ‘signature’ (i.e., the sinusoidal fluctuations in each signal channel) transfers from pitch to yaw in the orientation estimations. *Pitch*_*dp*_ is used to have a pose invariant descriptor of the gait of the animal, that is with respect to the animal itself rather than the environment. Inactive swimming periods (e.g., gliding) are automatically identified in *pitch*_*dp*_ while fluking frequency and amplitude are calculated from *pitch*_*dp*_, which can be used for further gait analysis.

## Discussion

The simulated datasets from bottlenose dolphins in a managed setting provided an efficient validation environment for the proposed methods. An extensive validation dataset that covered a wide variety of tag shifts was obtained. And, the ground truth measurements for the validation dataset were immediately available. The automatic shift detection algorithm is the first of its kind. This method demonstrated a detection precision of 0.87 and a recall of 0.89 with an average error of 37.5 seconds. The missed detections that were not ‘recalled’ by the method could be due to multiple injected shifts being temporally close to each other. As the method was designed to identify *occasional* tag shifts, the method may not detect all shifts if there are more than two shifts within two consecutive data segments. We determined the duration of a signal segment (*D*_*s*_) empirically to be 10 minutes (for bottlenose dolphins) to 15 minutes (for beluga whales and humpback whales). As demonstrated in Figure 8, if the duration of a data segment is too short, there is not enough data to form a meaningful orientation sphere for reliable shift detection. While if the duration of a data segment is too long, the method may not separately detect nearby shifts. It is recommended to use a duration *D*_*s*_ that is smaller than the expected tag shift interval, which could vary depending on the deployment. The situations being investigated in simulation were rather intense with relatively frequent tag shifts. For longer field deployments where the tag rarely shifts, the duration of a signal segment can be set to 20 minutes or longer to reduce false positives.

Another reason for missed detections could be that the tag shift was too small to be detected. The method’s sensitivity to shifting angles was investigated accordingly and presented in Figure 7. False positives (i.e., detections that did not correspond to a tag shift) were generated when the animals switched from one gait to another (very different) gait. For example, the method could be tricked when the animal changed from a gait without any roll to a sideways swimming gait (i.e., swimming with ∼90 degrees roll). In further developments, the method’s performance could be improved by taking into account events associated with high acceleration impacts to the tag, as tag shifts were often caused by physical impact to the tag [1], such as an impact from a conspecific animal in the group. Another aspect to be investigated in future development is the detection and management of consecutive shifts that happened within a period of time that can not be treated as *one* time instance (see footnote^5^). It is beyond the scope of this work to tackle the situation where the tag is constantly shifting.

To decouple the evaluations of tag shift detection and tag orientation correction, the tag orientation correction method was applied to each of the data segments transformed by a rotation. Poses (roll, pitch, and yaw) of the animal were calculated after orientation correction and then compared with poses computed using the original (un-rotated) data.

The average differences between the two were within 11 degrees in all cases (Table 1-**A** vs. **B** and Figure 7-Right). This result indicates that the tag orientation correction method can work reliably no matter where the tag was located as long as the given data segment did not contain a tag shift. One thing to note in particular is that the errors associated with pitch are higher than roll and yaw by a few degrees (Table 1-**A** vs. **B** and Figure 7-Right). This difference is likely because the tag was placed on the front half of the animal’s body between dorsal fin and blowhole (Figure 1-Left) in its ‘aligned’ position, which caused the ‘ground truth’ pitch measurement to have a negative bias. With this ‘aligned’ tag placement, the animal swimming in a horizontal direction would have a measured pitch centered around negative 6 ∼7 degrees instead of 0 degrees. In other words, the pitch measure returned after tag orientation correction could be more accurate than that obtained directly from an ‘aligned’ tag.

Meanwhile, the poses calculated using the stated method (involved accelerometer and magnetometer) in this work were compared with the established Madgwick’s filtering approach (involved accelerometer, magnetometer, and gyroscope [24], Table 1-**B** vs. **C**) and resulted in average differences that were within 3 degrees. The main error for the proposed pose estimation method came from estimating gravity direction using the accelerometer measurements. In this work, the low-pass filtered axis accelerometer data was used as the measurement associated with gravity (***A***^(*tag*)^) for the pose calculation. However, this is an inaccurate estimation in practice since the accelerometer measurements result from both gravity and the animal’s specific acceleration (i.e., the animal’s physical acceleration). The animal’s motion affects accelerometer measurements in 3 aspects:

(a1) Measurements associated with an animal’s specific acceleration. Since normally, the animal would not maintain a constant acceleration for more than a few seconds, all specific accelerations are considered high frequency.

(a2) Measurements associated with gravity, driven by **high** frequency body orientation changes.

(a3) Measurements associated with gravity, driven by **low** frequency body orientation changes.

Because measurements associated with gravity (a2 & a3) are needed for estimating animal pose, a1 needs to be decoupled from a2 and a3. An accurate decoupling would not be possible using an accelerometer alone. But because these animals’ specific accelerations are generally much smaller than gravity, a low-pass filter can be applied to attenuate a1 and a2 to approximately decouple specific acceleration measurement (a1) from gravity (a2 & a3).

To better decouple specific acceleration (a1) from gravity (a2 & a3), [33] proposed a *magnetometer* approach that takes advantage of the fact that a magnetometer is only sensitive to orientation changes, not acceleration. The method assumes that high-frequency body orientation changes are due to the ‘pitching’ motion of the animal. It relies on this assumption to use the magnetometer to assist in finding a2, with a3 obtained via low-pass filtering and a1 the remaining acceleration after subtracting a2 and a3. The method works well in decoupling a1 from a2 and a3 for most cases. However, we note that because the method assumes high-frequency body orientation changes in the pitching plane, the process tends to be unstable when the animal does quick back-and-forth rolling or turning. The method is implemented in the toolbox proposed in this work, and we recommend using it for animals with less frequent back-and-forth rolling or turning. For tags that have a gyroscope, together with an accelerometer and magnetometer, a gradient descent-based filtering approach [24] (*Madgwick* ‘s approach) can be used to better estimate the tag’s orientation wrt the world. The method uses the gyroscope to directly measure a2, thus reducing uncertainty in the measurement. A wrapper of Madgwick’s approach is also included in the toolbox, and we recommend using it when gyroscope measurements are available.

Even though the relative orientation of the tag was corrected to be aligned with the animal, the actual location of the tag (e.g., back vs. peduncle) could still affect tag measurements and results, particularly the estimated fluking amplitude of the animal. For example, a tag located closer to the fluke will have a higher estimated fluking amplitude than a tag located near the dorsal fin for the same gait. So, the fluking amplitude would be a good indicator of animal gait for fine-scale studies (e.g., comparing an animal’s fluking amplitude between descent vs. ascent), but not for global comparisons (e.g., comparing one animal to another, with different tag locations). On the other hand, the fluking frequency and period are not affected by the tag’s location and can be used for fine-scale studies and global investigations.

The orientation spheres (Figure 9) can also serve as a visualization tool for animal behavior studies. For example, a clear sign of lateralized behavior [34] appears in the orientation sphere of the humpback whale (Figure 9-Center Column), with an unbalanced right ‘wing’ corresponding to a dominating left roll of the animal during foraging at the ocean bottom. Also, this humpback whale did not roll much during ascending/descending while the bottlenose dolphin (Figure 9-Left Column) and beluga whale (Figure 9-Right Column) rolled more frequently during ascending/descending. Even though the beluga whale rolled frequently, it hardly ever rolled completely upside down, which can be observed from the empty bottom of the orientation sphere. Unlike the beluga, the dolphin data shown in Figure 9 indicate that the animal rolled all the way around during the studied period. In this regard, the presented orientation sphere, to a degree, resembles the *m-sphere* generated using magnetometer data in [35] as well as the *o-sphere* for visualizing animal’s head orientation in [31].

## Conclusion

This paper systematically presents an automated data processing framework (and software) that takes advantage of the common characteristics of pose and gait of the animal to process tag data and analyze the animals’ gait. Three aspects were investigated in particular: (1) Identifying time instances associated with occurrences of relative motion between tag and animal. (2) The identification of the relative orientation of a tag wrt the animal’s body for a given data segment. (3) The extraction of gait parameters that are invariant to pose and tag orientation. Biologging tag data from bottlenose dolphins, a humpback whale, and a beluga whale were used to validate and demonstrate the approach. Results show that the average relative orientation error of the tag wrt the dolphin’s body after processing were within 11 degrees in roll, pitch and yaw directions. In addition, the average precision and recall for identifying relative tag motion were 0.87 and 0.89, respectively. Examples of the resulting pose and gait analysis demonstrate the potential of this approach in gait analysis and animal behavioral studies. The proposed analysis approach and software will facilitate the use of biologging tags to study cetacean locomotion and behavior. The method and software is applicable to cetacean data from any tag platform that uses an accelerometer, magnetometer, and pressure sensor.

## Acknowledgments

We would like to thank Dolphin Quest Oahu for providing the scientific research environment and assistance during the experiments and for providing the in-kind support to make this project possible. The valuable discussions with Dr. Joaquin Gabaldon during the project are appreciated. We would also like to thank the funding sources that supported this work: the Canadian Department of Fisheries and Oceans (DFO), the US Office of Naval Research (ONR), and the US Navy’s Living Marine Resources (LMR) Program. The study protocols were approved by the University of Michigan Animal Welfare Committee (IACUC, #PRO00008825), the US National Marine Fisheries Service (NMFS, #18059), and the Canadian Council on Animal Care (#17-4, 18-3, 18-3B).

## Footnotes

1 *K* is an empirically determined constant value for the method. In this work, *K* = 30.

2 Near means the time interval is within an empirically defined threshold. In this work, the threshold interval is 3 minutes.

3 For the inertial coordinates, we define *x*^(*inertial*)^ points to magnetic north, *z*^(*inertial*)^ points vertically up, and *y*^(*inertial*)^ follows the right hand rule points to the magnetic west.

4 If ***V***_*dp*_ is applied to a shark, for example, the resulting vector would be mainly swinging in the 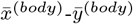 plane of the shark’s low frequency body coordinates.

5 If a sequence of consecutive shifts happened within a minute or two in a multiple-hour-long dataset, the sequence of shifts can be treated as one shift, per the resolution requirement of the application.

